# *Andropogon bicornis* L. (Poales: Poaceae): a stink bug shelter in the soybean and corn off-season in southern Brazil

**DOI:** 10.1101/675157

**Authors:** Eduardo Engel, Mauricio Paulo Batistella Pasini, João Fernando Zamberlan, Rafael Pivotto Bortolotto, Roberta Cattaneo Horn, Juliane Nicolodi Camera, Daniele Caroline Hörz

## Abstract

Associated host plants promote the survival of various species of pest insects during unfavorable periods. The objective of this study was to evaluate the diversity, abundance and structure of the pentatomid bugs community during the soybean and corn off-season in *Andropogon bicornis* L. (Poales, Poaceae) plants. The experiment was carried out in the municipality of Cruz Alta, RS, Brazil. During the soybean and corn off-season from 2014 to 2018, clumps of 10 to 50 centimeters in diameter were observed around the growing area. Data on the number of species and abundance of individuals were used for statistical analysis (ANOVA, linear regression and Pearson correlation) and faunistic (diversity and abundance distribution). At the end of the experiment 4050 adults belonging to the species *Euschistus heros* (F.), *Dichelops furcatus* (F.), *Dichelops melacanthus* (Dallas), *Edessa meditabunda* (F.), *Edessa ruformaginata* (De Geer) and *Piezodorus guildini*. Among the species, we found greater abundance for *E. meditabunda*, *E. heros* and *D. furcatus* (96.07%) of the individuals sampled. Higher interspecific correlation was observed between the same species, we also observed a direct effect of the clump diameter on the population density. Among the six species observed, at least five are economically important for soybean and corn crops, so further studies are needed in order to verify the effects of this hibernation site on the population present in the crop and its damages.

## 1. Introduction

*Andropogon bicornis* L. plants (Poales: Poaceae) are organisms of significant abundance in the south of Brazil, being distributed perennially throughout the landscapes, constituting important part of the phytogeography of this region (Boldrini, 2009; Santos et al. 2015). *A. bicornis* is distributed along the borders of roads and borders of crops cultivated with diverse crops, with soybeans and corn being important crops surrounded by these plants, in this way, this plant may be associated to several arthropods occurring during the culture in both cultures (Kissmann & Goth, 2000; Klein, Redaelli & Barcellos, 2012; Pasini, Lúcio & Ribeiro, 2015).

Several works demonstrate the importance of the plant structures surrounding the cropping areas in the survival of pest insects throughout the year, this importance is related mainly to the periods of low temperatures and availability of food, in this way, they constitute adequate shelters for hibernation (Dennis, Thomas & Sotherton, 1994; Klein, Redaelli & Barcellos, 2012, 2013; Botta et al., 2014; Pasini, Lúcio & Ribeiro, 2015; Pasini et al., 2018).

Portraits of the Pentatomidae family are among the main entomological problems for diverse cultures worldwide (McPherson, 2018) their damages and feeding behavior were extensively studied, thus confirming their economic importance for soybean and corn (Cooke, 2014; Lucini & Panizzi, 2017 a, b, c). Knowing the behavior of these individuals in natural landscapes constitutes an important tool for decision making in Integrated Pest Management (IPM) (Mendonça, Schwertner & Grazia, 2009).

Therefore, the objective of this study was to verify the diversity, structure and abundance of stink bugs associated to *A. bicornis* in the soybean and corn intercrop for five years.

## 2. Method

The experiment was conducted during the month of June 2014 to 2018 at the Experimental Area at the University of Cruz Alta (Time Zone 22, 244138; 6835737). According to Köppen, the climate of the study area belongs to type Cfa, with average temperature below 18°C in the coldest month and average temperature above 22 ° C in the hottest month of the year (Kuinchtner & Buriol, 2016).

Perennial plants of *Andropogon bicornis* L. (Poales: Poaceae) were sampled. The plants sampled were located at a distance of 20 meters from the edge of the growing area, between the plants, the minimum distance adopted was 15 meters. Fifty specimens were sampled per year, divided into five different clump diameters (0-10, 10-20, 20-30, 30-40 and 40-50 centimeters). Each plant was considered an experimental unit (EU), totalizing at the end of the experiment 250 EU.

The stink bugs sampled in each plant were identified and counted for later data analysis. For those not identified, these were separated into morphospecies and taken to the Laboratory of Entomology - UNICRUZ for identification and accounting.

In order to analyze the population density of stink bugs in the evaluated plants, the data were submitted to the normality and homogeneity tests of Anderson-Darling and Bartlet variances respectively. For the data that did not meet the assumptions of the tests, these were submitted to function Root (x+0.5). After normalization, the data were submitted to ANOVA, and the means were compared by the Tukey test at 5% of error probability. To determine the relationship between clump diameter and number of stink bugs, we used linear regression analysis. To determine the correlation between the occurring species, the data were analyzed by Pearson’s linear correlation with a 5% probability of error.

After identification of the occurring species, the quantitative data were used for analysis through alpha diversity (species diversity, distribution of species abundances - SAD). The suitability of SAD was tested in four models: geometric, broken-stick, log-series and log-normal. The sampling adequacy curve for the abundance of *A. bicornis* associated pentatomids was obtained with 999 randomizations and compared with the non-parametric wealth estimators Chao 2, Jacknife 1 and Jacknife 2 to determine the sampling efficiency, according to the methodology used by Bianchi et al. al. (2018). Each wealth estimator takes into account a different parameter, that is, occurrence of singletons, doubletons, unique and duplicates. All analyzes were performed using the PASt 3.18 software (Hammer, Harpista & Ryan, 2001).

## 3. Results and Discussion

At the end of the experiment, the number of stink bugs sampled was 4050 individuals distributed in six species (Table 1).

**Table 1.**
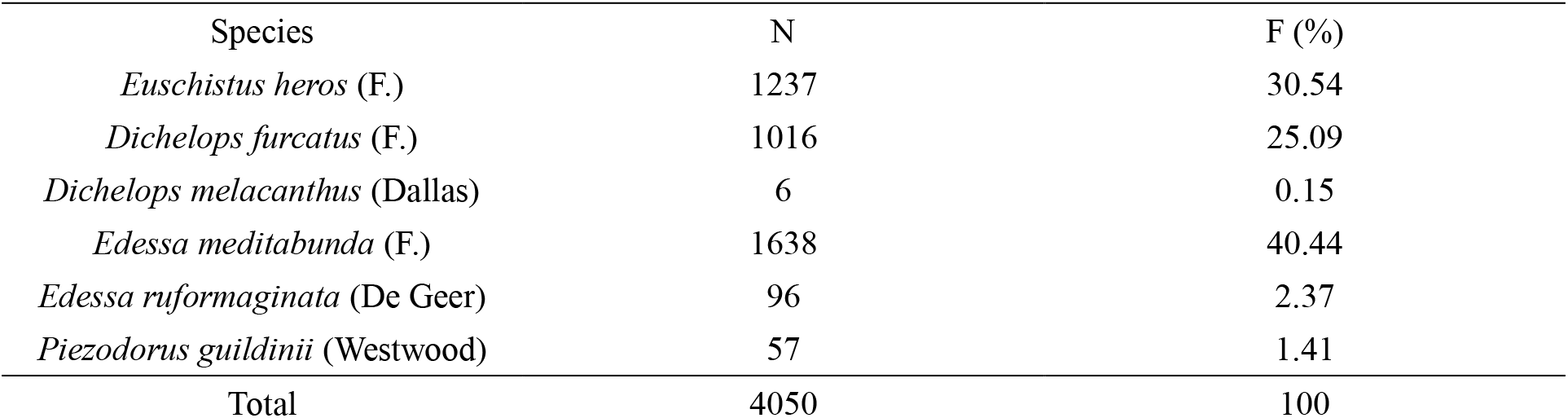
Species, abundance (N) and frequency (F) of stink bugs sampled in *Andropogon bicornis* L. (Poales: Poaceae) in the soybean and corn off-season.

Among the species, we observed a high frequency of *E. meditabunda* (40.44% of the individuals sampled), followed by *E. heros* and *D. furcatus* (30.54 and 25.09%, respectively). *E. meditabunda* for Brazil is considered a secondary pest for soybean crop, feeding on vegetative parts of the crop (stems and leaves), but in Argentina this insect has high economic importance and is considered a key pest (Saluso et al 2007, Panizzi & Silva 2012). However, *E. heros* and *D. furcatus* have high economic importance for soybeans and corn in Brazil, are considered highly polyphagous insects, having a high number of host plants, in this way, are able to remain in the area of cultivation and its proximity for long periods (Panizzi, Bueno & Silva, 2012; Smaniotto & Panizzi, 2015).

A tendency to stabilize the number of bed bug species was observed in the rarefaction curve, with the observed values (6) being very close to or equal to the estimated values (Chao 2: 6.01, Jackniffe 1: 5.99, Jackniffe 2: 6.00 and Bootstrap: 6.04) (Figure 1). These values indicate sufficiency in the number of samples employed, contemplating all the species of pentatomid bugs that occur in this plant in the study area. In the rice agroecosystem, Klein, Redaelli & Barcellos (2013) verified a greater species richness, contemplating 14 species of pentatomid bugs, being the rarefaction curve indicating probability of a greater number of occurring species.

**Figure 1.**
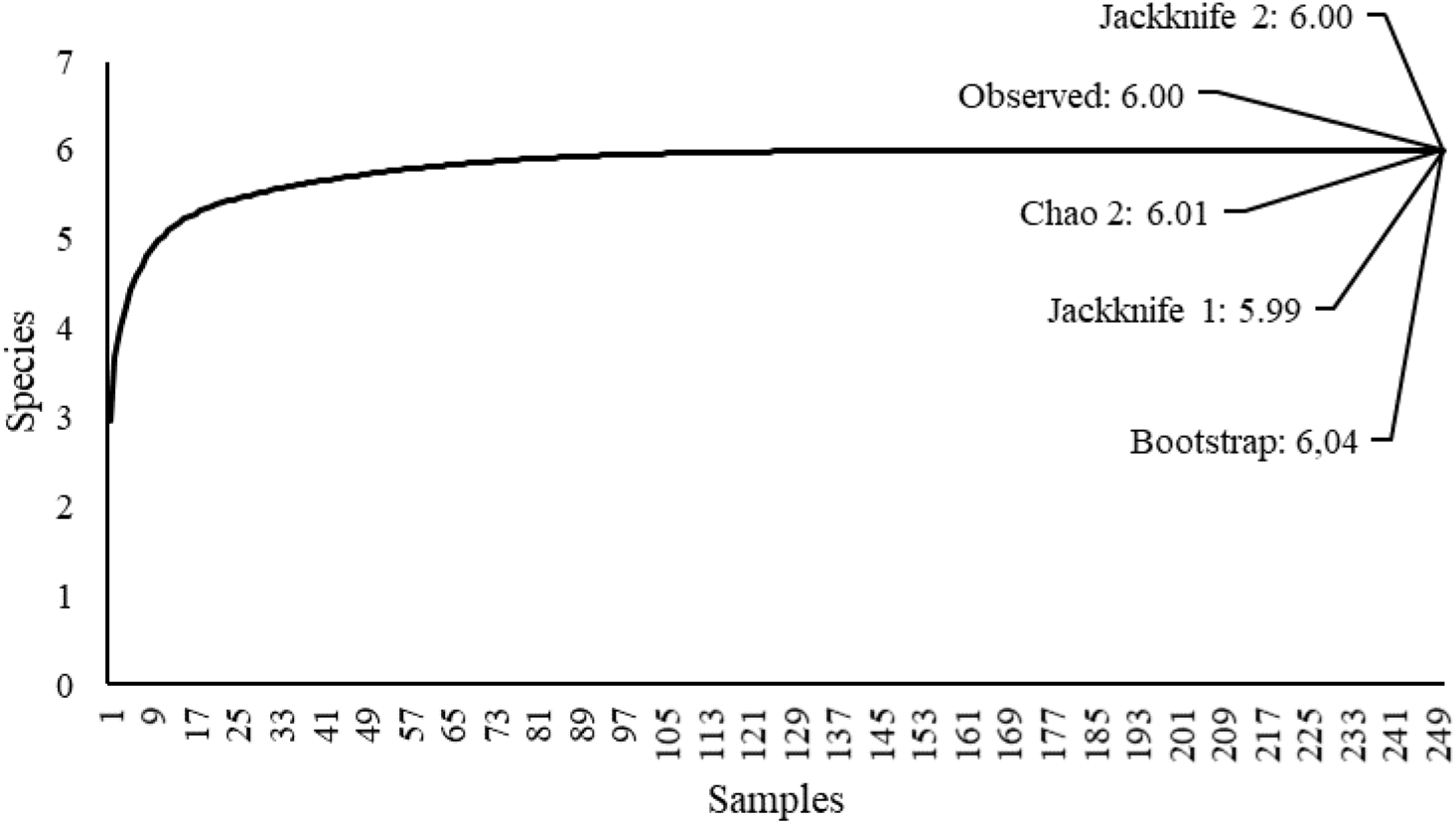
Rarefaction curve and species richness estimators for pentatomid bugs sampled in *Andropogon bicornis* L. (Poales: Poaceae) during the soybean and corn off-season from the years 2014 to 2018. Cruz Alta, Rio Grande do Sul, Brazil.

Among the species, we found that *E. meditabunda* was significantly more abundant, followed by *E. heros* and *D. furcatus* (Figure 2). These three species constitute 96.07% of all individuals sampled. The SAD analysis showed significance for the geometric model (K = 0.6778; Chi ^ 2 = 2192; p = 0.00 <0.05) (Figure 3). In this model, the species with the greatest abundance is responsible for the use of most of the available resources, followed by the other species respectively (Magurran, 2004; Bianchi et al., 2018).

**Figure 2.**
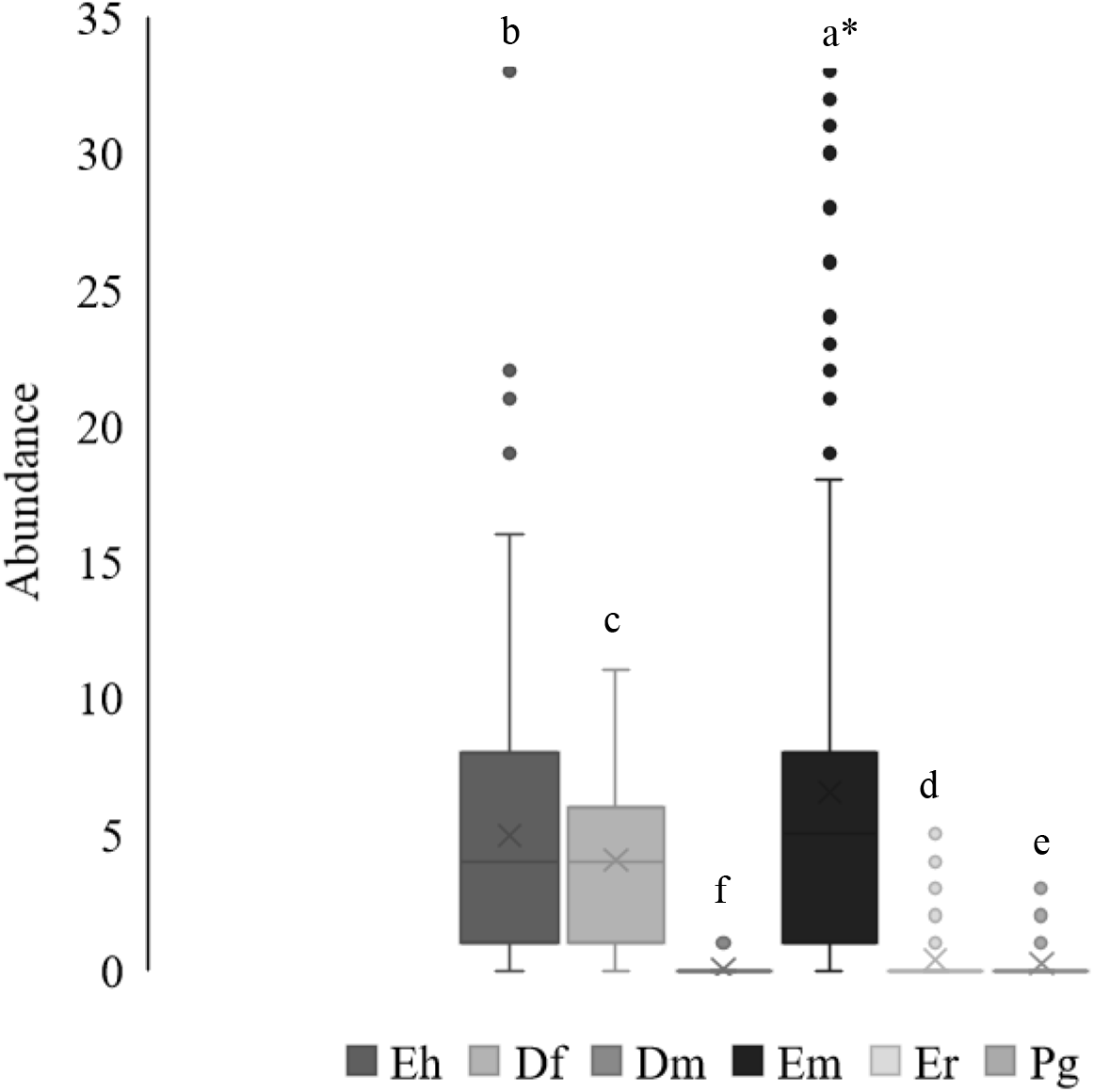
Box plot of the number of stink bugs sampled in *Andropogon bicornis* L. (Poales: Poacaeae) during the soybean and corn off-season from the years 2014 to 2018. Cruz Alta, Rio Grande do Sul, Brazil. Eh (*Euschistus heros*), Df (Dichelops furcatus), Dm (Dichelops melacanthus), Em (Edessa meditabunda), Er (*Edessa ruformaginata*) and Pg (*Piezodorus guildinii*). * Letters differ from each other by the Tukey test (p<0.05).

**Figure 3.**
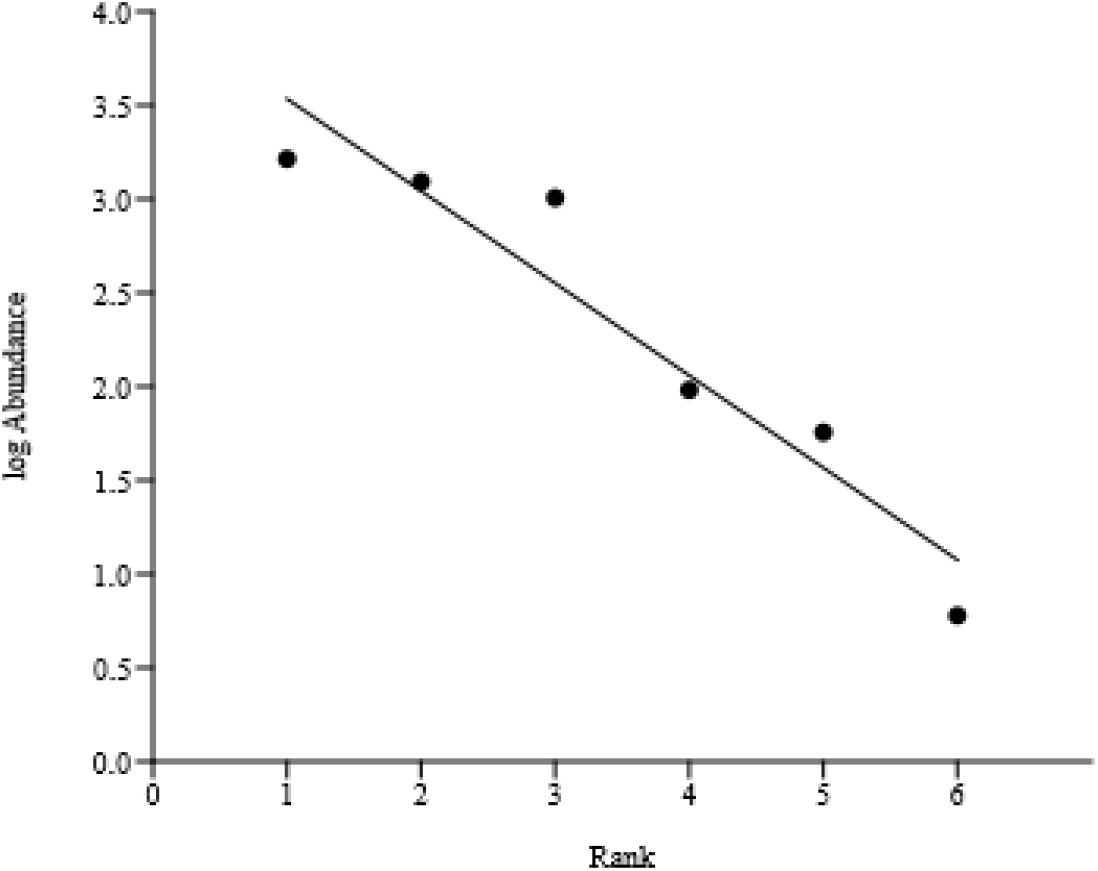
Distribution of the abundances of pentatomid bugs sampled in *Andropogon bicornis* L. (Poales: Poaceae) in the soybean and corn off-season al from the years 2014 to 2018. Cruz Alta, Rio Grande do Sul, Brazil.

We verified a significant correlation (p <0.05) among the species, being the highest among E. heros *and D. furcatus* and *E. meditabunda* and *E. heros* (Fig 4). For other species we verified a lower correlation index. This may be associated with low population density for *D. melacanthus* and *P. guildinii*, indicating no preference for hibernation in *A. bicornis* plants. *E. ruformaginata* presents different behavior of the other species, having a higher occurrence in wild vegetation, being not related to soybean and corn crops (Silva & Oliveira, 2010).

**Figure 4.**
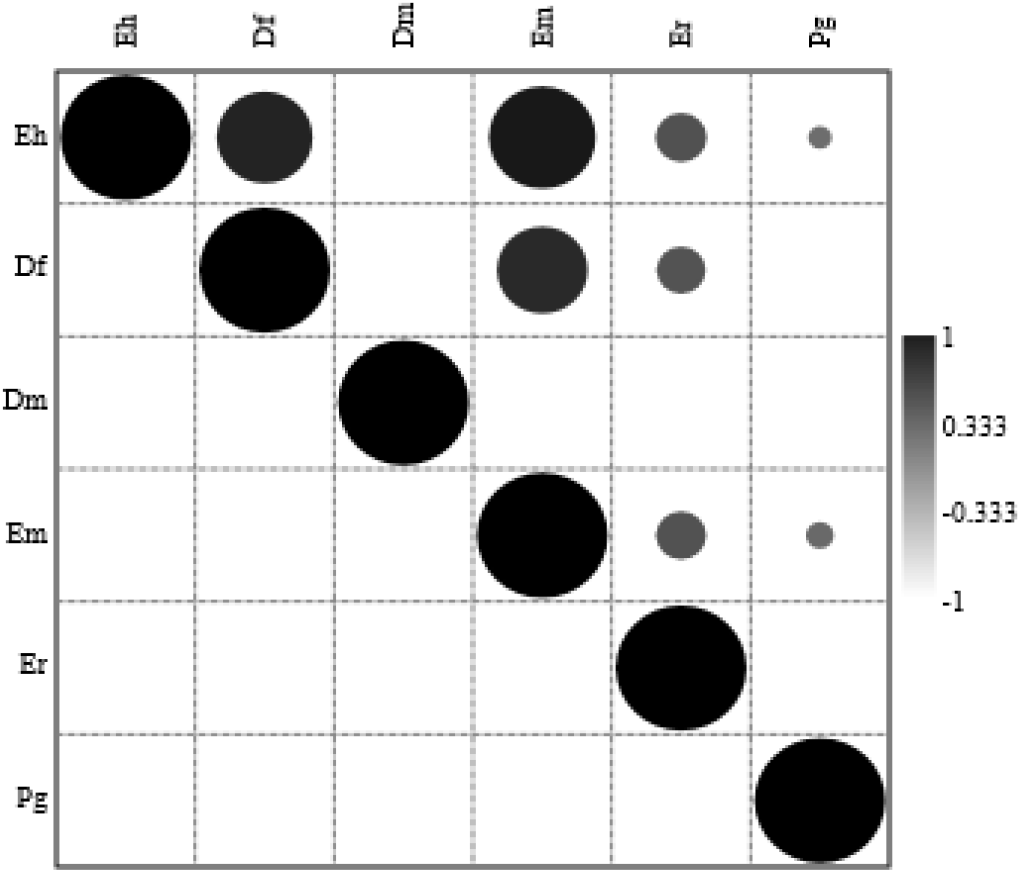
Pearson’s linear correlation for stink bugs occurring in *Andropogon bicornis* L. (Poales: Poaceae) plants in the soybean and corn off-season from 2014 to 2018. Cruz Alta, Rio Grande do Sul, Brazil. Eh (*Euschistus heros*), Df (*Dichelops furcatus*), Dm (*Dichelops melacanthus*), Em (*Edessa meditabunda*), Er (*Edessa ruformaginata*) and Pg (*Piezodorus guildinii*).

Another factor of significant importance was the direct relationship between the diameter of the clumps evaluated and the population density, we observed an abrupt increase in the density of stink bugs due to increase of the clump diameter (Figure 5). Para Klein, Redaelli & Barcellos (2013) clumps with larger structures provide less space dispute and more stable microclimate conditions than in open environments. Several studies demonstrate the importance of plant structure in stink bug sampling, where plants with a higher level of complexity and volume tend to offer greater opportunities for survival of these organisms (Howe & Jander, 2008; Klein, Redaelli & Barcellos, 2013; Knolhoff & Heckel, 2014; Pasini, Lúcio & Ribeiro,2015; Pasini et al., 2018).

**Figure 5.**
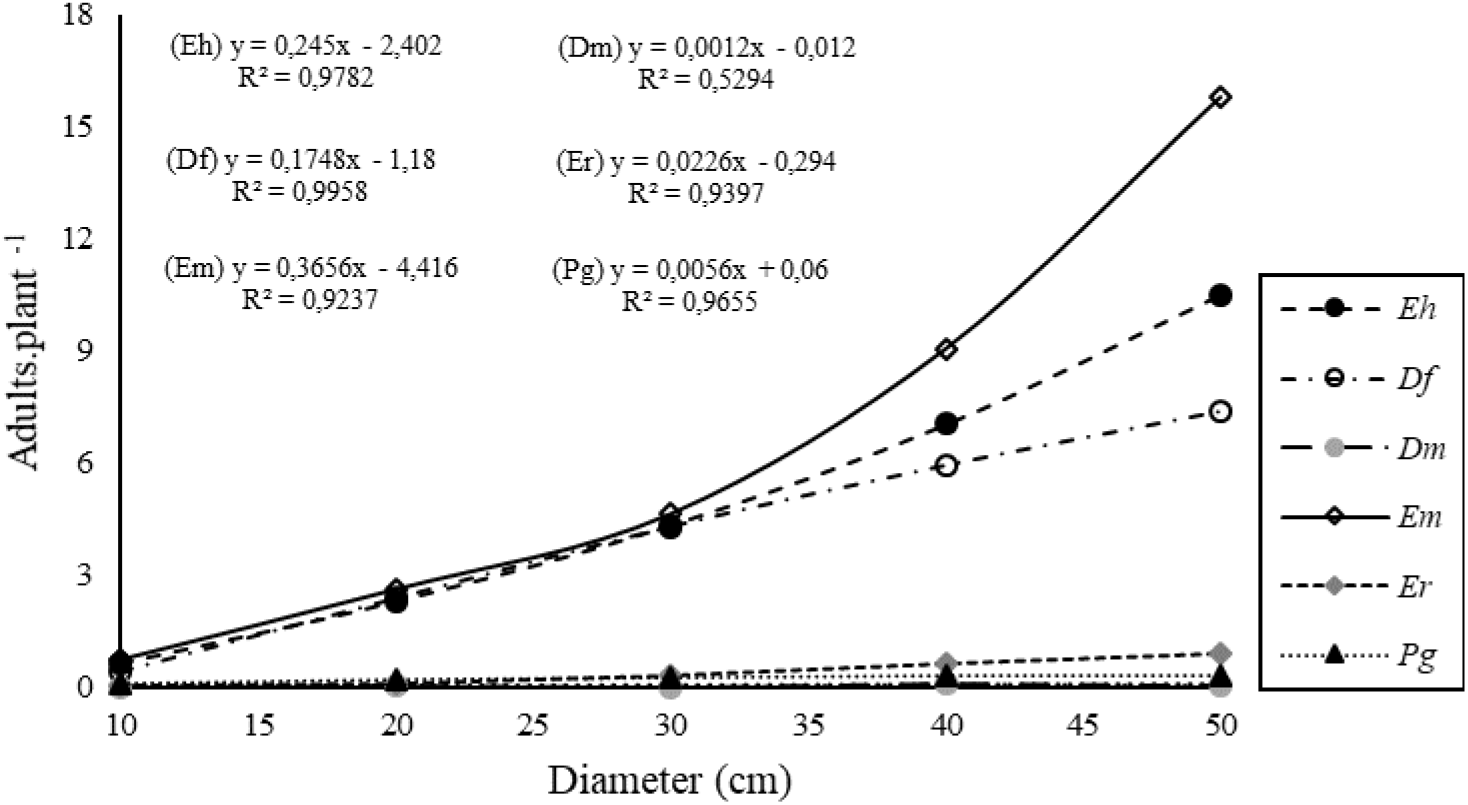
Population density of phytophagous bugs as a function of host plant diameter. Eh (*Euschistus heros*); Df (*Dichelops furcatus*); Dm (*Dichelops melacanthus*); Em (*Edessa meditabunda*); Er (*Edessa ruformaginata*); Pg (*Piezodorus guildinii*). Cruz Alta, Rio Grande do Sul, Brazil, 2014 to 2018.

The presence of *A. bicornis* in the areas surrounding the crops adds greater diversity in the landscape, this directly influences the diversity of arthropods (Helden et al., 2010; Klein, Redaelli & Barcellos 2013). As in the rice landscape, *A. bicornis* is also used as a shelter for several species of stink bugs of economic importance in the soybean-corn scenario, being an important source of reinfestation for the crops, as well as a weed for both crops. Their clumps directly influence the sheltered population density. The presence of nymphs and eggs were not verified in any of the evaluated years, indicating the use of this plant for hibernation purposes only. Further studies should be performed to understand the impact of their management on the population of stink bugs and their damage.

## 5. Conclusions

The study evidenced the importance that A. bicornis has for the survival of stink bug species considered important pests for soybean and corn crops. Its morphological structure directly interferes with population density. We verified that *E. meditabunda*, *E. heros* and *D. furcatus* presented the largest populations Further studies should be carried out in order to determine the effect of the management of these host plants on the population density and damages of these stink bugs.

